# Biomarker2vec: Attribute- and Behavior-driven Representation for Multi-type Relationship Prediction between Various Biomarkers

**DOI:** 10.1101/849760

**Authors:** Zhen-Hao Guo, Zhu-Hong You, Yan-Bin Wang, Hai-Cheng Yi

## Abstract

The explosive growth of genomic, chemical and pathological data provides new opportunities and challenges to re-recognize life activities within human cells. However, there exist few computational models that aggregate various biomarkers to comprehensively reveal the physical and functional landscape of the biology system. Here, we construct a graph called Molecular Association Network (MAN) and a representation method called Biomarker2vec. Specifically, MAN is a heterogeneous attribute network consists of 18 kinds of edges (relationships) among 8 kinds of nodes (biomarkers). Biomarker2vec is an algorithm that represents the nodes as vectors by integrating biomarker attribute and behavior. After the biomarkers are described as vectors, random forest classifier is applied to carry out the prediction task. Our approach achieved promising performance on 18 relationships, with AUC of 0.9608 and AUPR of 0.9572. We also empirically explored the contribution of attribute and behavior feature of biomarkers to the results. In addition, a drug-disease association prediction case study was performed to validate our method’s ability on a specific object. These results strongly prove that MAN is a network with rich topological and biological information and Biomarker2vec can indeed adequately characterize biomarkers. Generally, our method can achieve simultaneous prediction of both single-type and multi-type relationships, which bring beneficial inspiration to relevant scholars and expand the medical research paradigm.

## Introduction

A key task in the post-genomic era is to systematically and comprehensively understand the relationships between molecules in living cells [1]. Rapidly developing high-throughput technologies and the discoveries of new transcripts or translations provide foundation for this mission [2]. For example, the increasing evidence prove that the biology molecule networks such as protein-protein interaction network, ncRNA-disease association network, drug-target interaction network play significant roles in protein synthesis [3], gene expression [4], RNA processing [5] and developmental regulation [6], etc. Consequently, microscopic research of the relationships between biomarkers not only opens novel insights to understand life process, but also facilitates to disease prevention, diagnosis, treatment and drug development.

Identifying the relationships in large-scale data via wet experiments is labor-intensive, time-consuming and can only meet limited requirements in real-world demands. Meanwhile, the extensive accumulated experiment data lead to the trouble of information overload and the excess cost of acquiring valuable knowledge. Hence, it is urgent to design automatic computational tool to provide assistance and guidance for practice [7].

In fact, prediction models based on validated evidence to discover potential relationships have been widely developed and heavily applied. Guo *et al.* proposed a learning-based model to predict potential lncRNA-disease associations by integrating known association evidence, disease semantic similarity [8]. Wang *et al.* carry out the Logistic Model Tree to discover unknown miRNA-disease associations by integrating multi-source information [9]. Li *et al.* used the Position-Specific Scoring Matrix (PSSM) to represent proteins and then put them into an ensemble classifier to predict self-interacting and non-self-interacting proteins [10]. Wang *et al.* utilize the Rotation Forest as a classifier to uncover unknown drug-target interactions by drug structure and protein sequence [11].

Although many attempts have been made to detect the uncovered relationships through various methods including matrix factorization [12], machine learning [13] and network analysis [14]. The incompleteness of the data constrains the credibility of the prediction results accompanied with higher FPR and FNR [15]. In recent years, the discovery of new types biomolecules and relationships provides novel insights to improve this situation to some extent. Biomolecules outside the research subject are attempted to be considered as bridges, and synergistically help the grasp of underlying biological principles to improve the prediction effect. Chen *et al.* deeply understand the pathogenesis of disease by environmental factors and effectively improve the prediction effect of miRNA-disease association [16]. Cui *et al.* make a preliminary exploration in the prediction of drug-disease drawn support from gene expression data [17].

Tremendous advances in molecular biology over the past few years, yet the development of computational model is still in infancy. The major limitation of all above methods is that none of them regard a cell as a complete unit, even if these methods have their own unique advantages. In fact, cells are composed of nodes (biomarkers) and edges (relationships) like a network (graph) to maintain normal life activities and physiological functions. Ideally, establishing connections from internal or external factors to expression would be rewarding in understanding the landscape of the biology system. In this paper, a complete network called Molecular Association Network (MAN) is constructed based on various online database such as NONCODE [18] and miRbase [19] to provide a platform to help systematically analyze the inseparable connections and messages flow between biomarkers within human cells.

Faced with such a large-scale network, the most critical challenge is how to quickly and effectively describe the relationships between nodes. The rapid development of graph embedding (network representation) algorithms has shown us hope for addressing such problems [20]. Graph embedding which aims to represent nodes in the network as low-dimensional, dense vector forms is chosen to respond to this situation [21]. Although some existing models in bioinformatics contain the idea of graph embedding, many of them still focus on traditional techniques including Principal Component Analysis (PCA) [22], Multidimensional scaling (MDS) [23], Isomap [24] and Local Linear Embeddings (LLE) [25]. In general, these methods offer satisfactory performance on small networks. However, at least quadratic time complexity restricts the application of these methods to large-scale data. The recent remarkable performance of deep learning has attracted extensive research attention. Here, the representation method called DeepWalk is applied to conduct this task.

Attribute feature including RNA sequence, drug structure and disease semantics is another available information and the potential of them in inferring potential relationships has been widely documented. Frequently in these methods, the attribute of each biomolecule is represented as a vector using various feature representation methods.

In this paper, a network called Molecular Association Network (MAN) is constructed and a graph embedding algorithm is proposed to represent each node as a vector. Then random forest is applied as the classifier to carry out the relationship prediction task. Specifically, 18 kinds of associations or interactions among 8 kinds of biomolecules are collected from various database to construct the Molecular Association Network (MAN). Then we develop the lower triangular part of the adjacency matrix called *A* containing the whole information of the graph to simplify calculation and storage. Obviously, each node in the network can be described from 2 perspectives as shown in the Figure 1, one is the attribute feature such as sequence and chemical structure that can be learned as a 64-dimension vector by k-mer and etc. methods, the other is the behavior feature that is the relationships that can be represented as a 64-dimension vector through DeepWalk. Stack autoencoder is applied to unify the dimension and improve feature quality. Then each node can be represented as a 128-dimension vector by Biomarker2vec through integrating attribute and behavior feature. The positive samples are experimentally verified relationships and the negative samples are the same number of unlabeled relationships which are randomly selected in *A*. Taking the low-dimensional dense vectors as input, random forest is used to carry out the prediction task. The proposed method obtained AUC of 0.9608 and AUPR of 0.9572 under 5-fold cross validation on the multi-type relationship prediction task of whole network. Furthermore, we implemented 3 comparison experiments including feature importance comparison, embedding strategy comparison and proportions of training set comparison. The remarkable performance demonstrated that MAN with bright prospects of revealing uncovered relationships in human cells. We hope that this work can provide assistance and guidance for wet experiments, and be a useful inspiration for researchers to understand gene regulation, disease mechanism and discovery of new drugs at molecular level.

**Figure 1.**
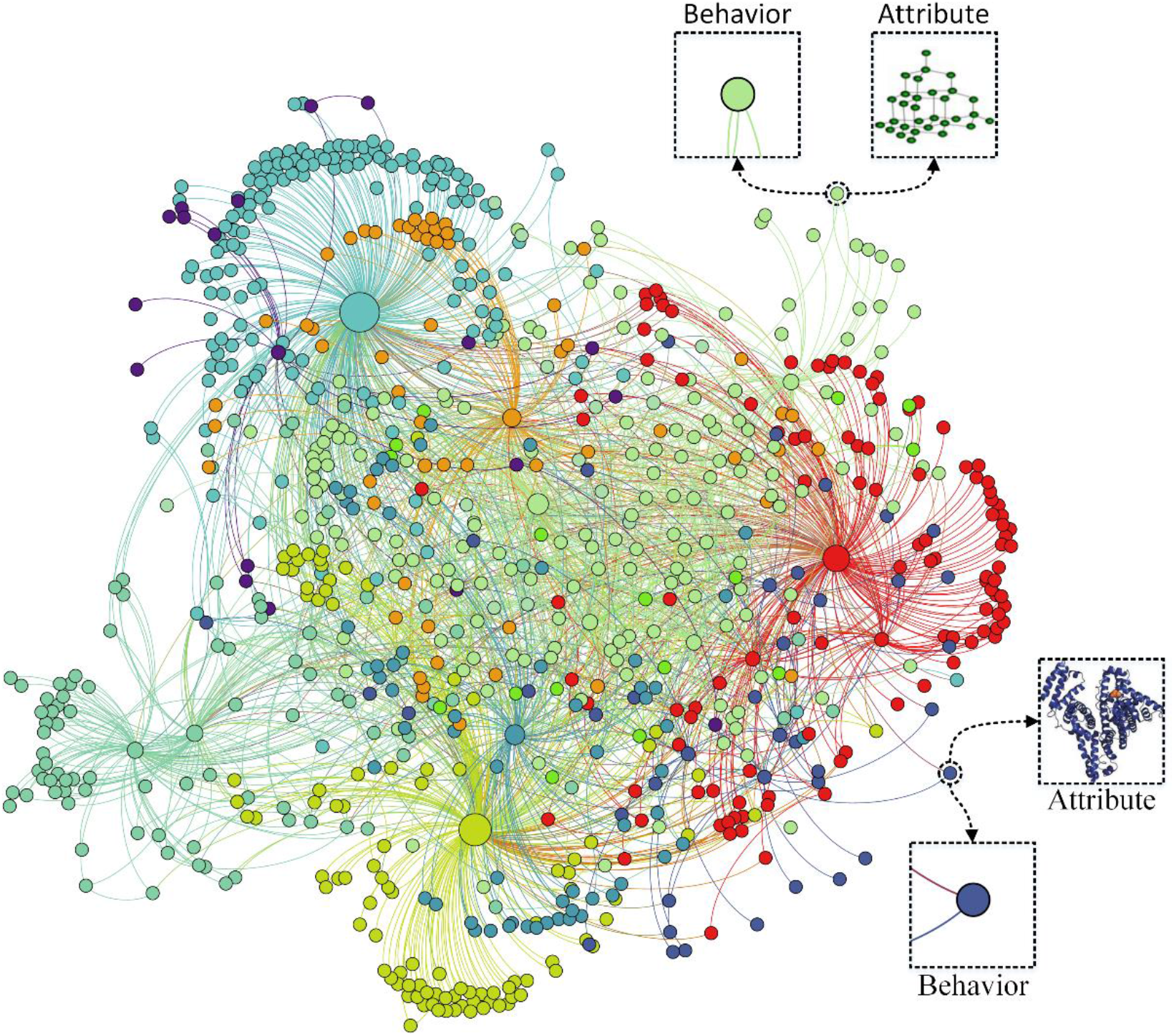
A visualization example like MAN, where different colors represent different types of biomarkers. Each biomarker contains 2 kinds of information including node behavior (relationships with other nodes) and node attribute (sequences of protein or RNA, chemical structure of drug, and semantics of disease and microbe).

**Figure 2.**
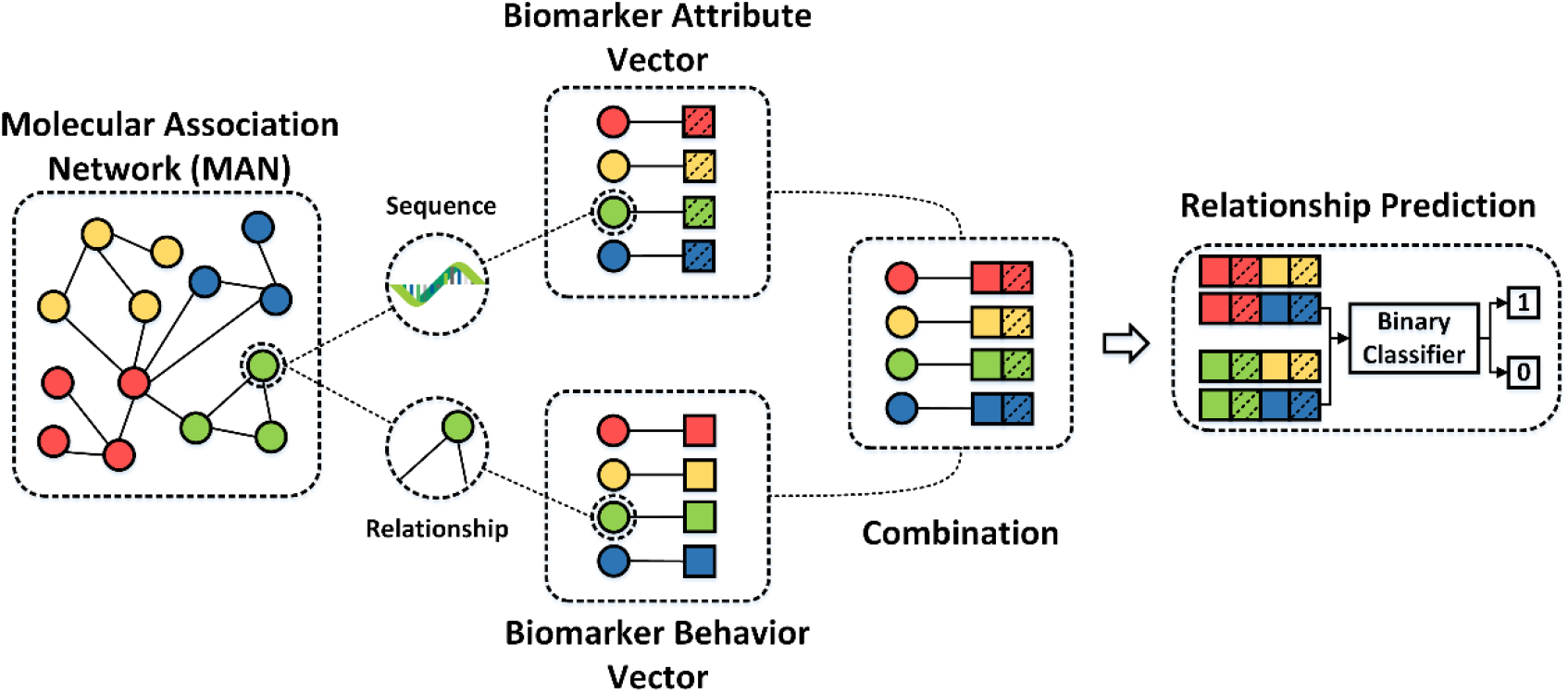
The flowchart of proposed method.

## Materials and Methods

### 2.1 Construction of the Molecular Association Network (MAN)

To construct the Molecular Association Network (MAN) comprehensively, 18 different kinds of experimental verified associations or interactions are collected from various databases [26–45]. After the unifying identifier, we obtained a total of 8 diverse types of biomarkers. Then, all relationships and biomarkers are aggregated together to form the complete MAN. The specific quantity and proportion of each type of nodes or relationships are shown in the figure below.

**Figure 3.**
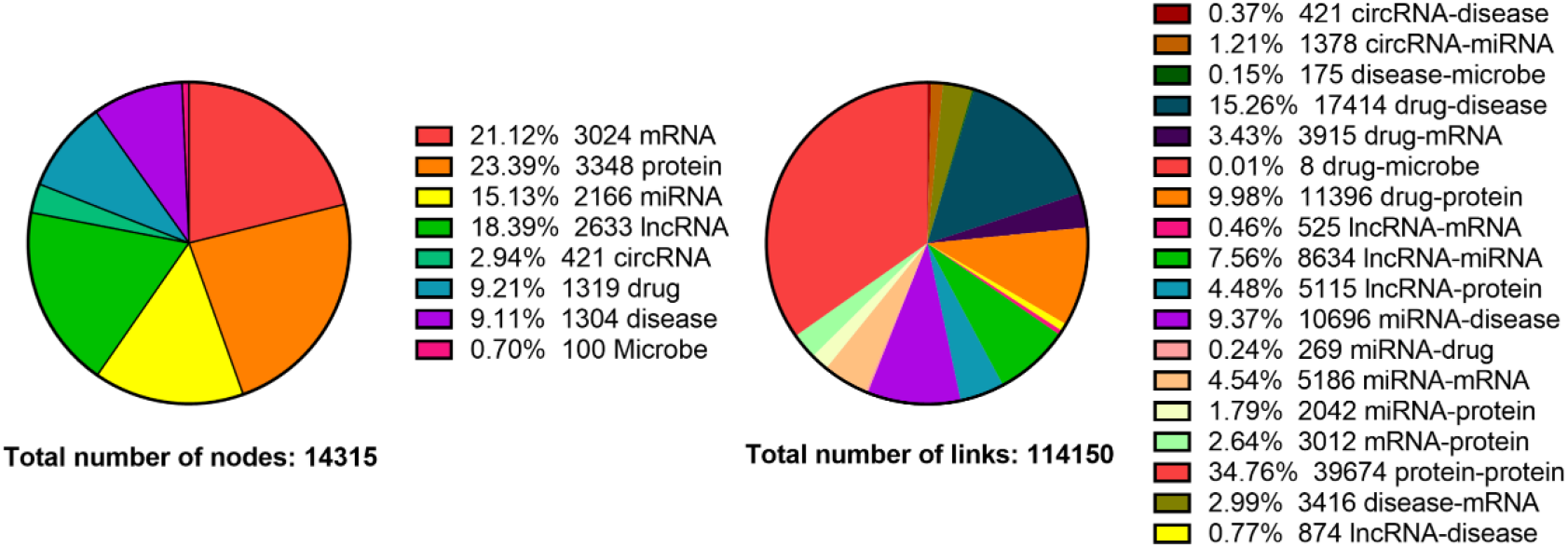
The details of different biomolecules and relationships

### 2.2 Node attribute representation: K-mer, Semantics and Fingerprint

It is obvious that the intrinsic attribute such as the sequence of protein and RNA, the semantics of disease and microbe, and the chemical structure of drug is the essential feature of each biomolecule. The details of how they are represented as vectors are as follows.

For protein, mRNA, miRNA, lncRNA and circRNA, their sequences are collected from STRING [44], NCBI, miRBase [19], NONCODE [18] and circBase [46] respectively. Given that proteins are composed of 20 different types of amino acids and inspired by the method of Shen *et al.* [47], we first classify them into 4 categories based on the polarity of the amino acid side chains including (Ala, Val, Leu, Ile, Met, Phe, Trp, Pro), (Gly, Ser, Thr, Cys, Asn, Gln, Tyr), (Arg, Lys, His) and (Asp, Glu). RNA including mRNA, miRNA, lncRNA and circRNA are composed of 4 kinds of nucleotides including Adenine (A), Guanine (G), Cytosine (C) and Uracil (U) with the same sequence composition, so we directly encode their original sequence without any pretreatment. Each RNA or Protein can be represented as a vector by k-mer, in which all dimensions represent the full permutation of k nucleotide combinations and the value of each dimension is the normalized frequency of the corresponding k-mer appearing in the sequence. In this paper, k is set to 3 and each protein or RNA can be represented as a 64-dimension (4^3^ = 4 × 4 × 4) vector.

For disease and microbe, their Medical Subject Headings (MeSH) descriptors which are comprehensive control vocabularies organized by U.S. National Library of Medicine are downloaded from https://www.nlm.nih.gov/. The top-level categories in the MeSH Tree Structure are: Anatomy [A], Organisms [B], Diseases [C] and so on. The categories corresponding to microbes and diseases are B and C, respectively. Inspired by D Wang *et al.* [48], we construct the Directed Acyclic Graph (DAG) of disease and microbe to represent them through their semantics. For example, a microbe *M* can be represented as a graph *DAG*(*M*) = (*M*, *N*(*M*), *E*(*M*)) where *N*(*M*) is the set of all nodes in *M*’s DAG and *E*(*M*) is the set of all edges in *M*’s DAG. The semantic contribution of microbe *m* which is in the node set *N(M)* to *M* can be defined as:

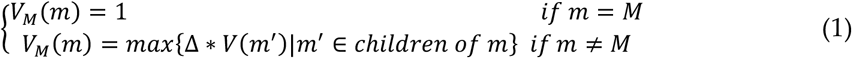

where Δ denotes an attenuation factor and is defined as 0.5 according previous literature. In the DAG generated by microbe *M*, *M*’s contribution to itself can be regarded as the maximum and equals to 1, and the remaining diseases will contribute less and less to *M* as the distance increases. Therefore, the sum of the contributions of microbes which are in the set *N(M)* to *M* can be calculated as follows:

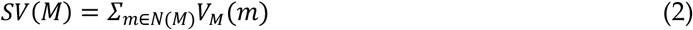

Then the similarity between microbe *i* and *j* can be calculated by the following formula:

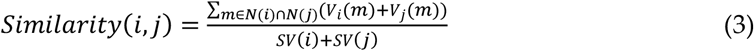

The node attribute of microbe or disease can be represented by semantics similarity, which is converted into a 64-dimensional vector after feature extraction and transformation by the stack autoencoder. A DAG example of microbe Staphylococcus is as follows:

For drug, we download their SMILES [49] from DrugBank [35] and transform the SMILES into corresponding Morgan Molecular Fingerprints [50] by python package called RDKit [51]. To unify dimensions and improve feature quality, stack autoencoder is used to convert each original molecular fingerprint into a 64-dimensional vector.

**Figure 4.**
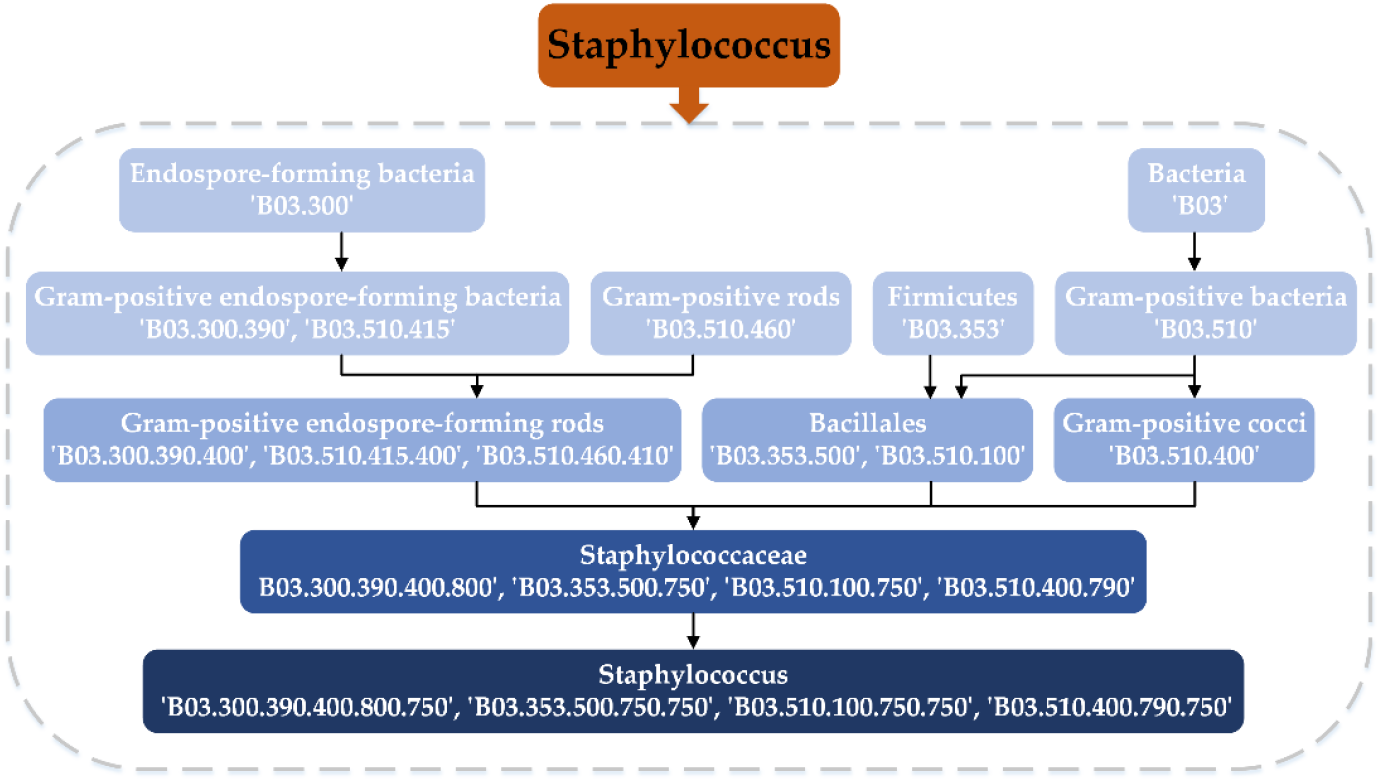
Construction the DAG of Staphylococcus. The father node of the current microbe can be obtained by deleting the last three digits of the descriptor. For example, for Bacillales (B03.353.500, B03.510.100), we can remove the last three digits to get Firmicutes (B03.353) and Gram-positive bacteria (B03.510).

### 2.3 Node behavior representation: DeepWalk

Inspired by the idea of “guilt-by-association” assumption, we come up with a more general feature in complex networks, that is, the behavior feature of biomarker. Generally speaking, it is a kind of embedding representation of the known edges between nodes in the network. Efficient description of the behavior feature is the core issue of relationship prediction on large-scale biomolecule networks. Despite a row or column of the adjacency matrix can directly be utilized as a representation vector for node behavior in one-hot encoding method. However, there is no concept of similarity between each dimension of such high-dimensional, sparse vectors, as it is represented as indices in a relationship. Meanwhile, the one-hot encoding method takes up a lot of storage space and is not conducive to the input of downstream tasks. Hence, how to extract the behavior information of node from the complex network such as MAN is a formidable challenge.

In this paper, a network embedding method called DeepWalk which first applies the technique of natural language processing in deep learning for node representation is adopted to undertake this task [52]. The main idea is to obtain a certain length of the walk sequence through RandomWalk, an ideal mathematical state of Brownian motion that can repeatedly access the visited nodes. After obtaining enough sequences, the vectors of the nodes can be learned by the skip-gram model. The direct analog is to estimate the likelihood of observing vertex *v*_*i*_ given all the previous vertices visited so far in the random walk, *i.e.*

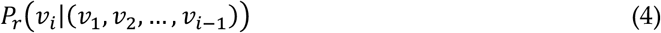

The goal is to learn a latent representation and the mapping function is:

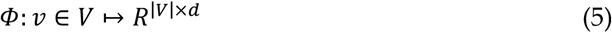

The problem then, is to estimate the likelihood:

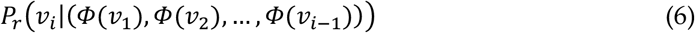

The recent relaxation in language modeling turns the prediction problem and this yields the optimization problem:

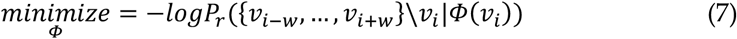

The main steps of the algorithm are as follows:

#### Algorithm 1

DeepWalk (***G***,***w***,***d***,***γ***,***t***).

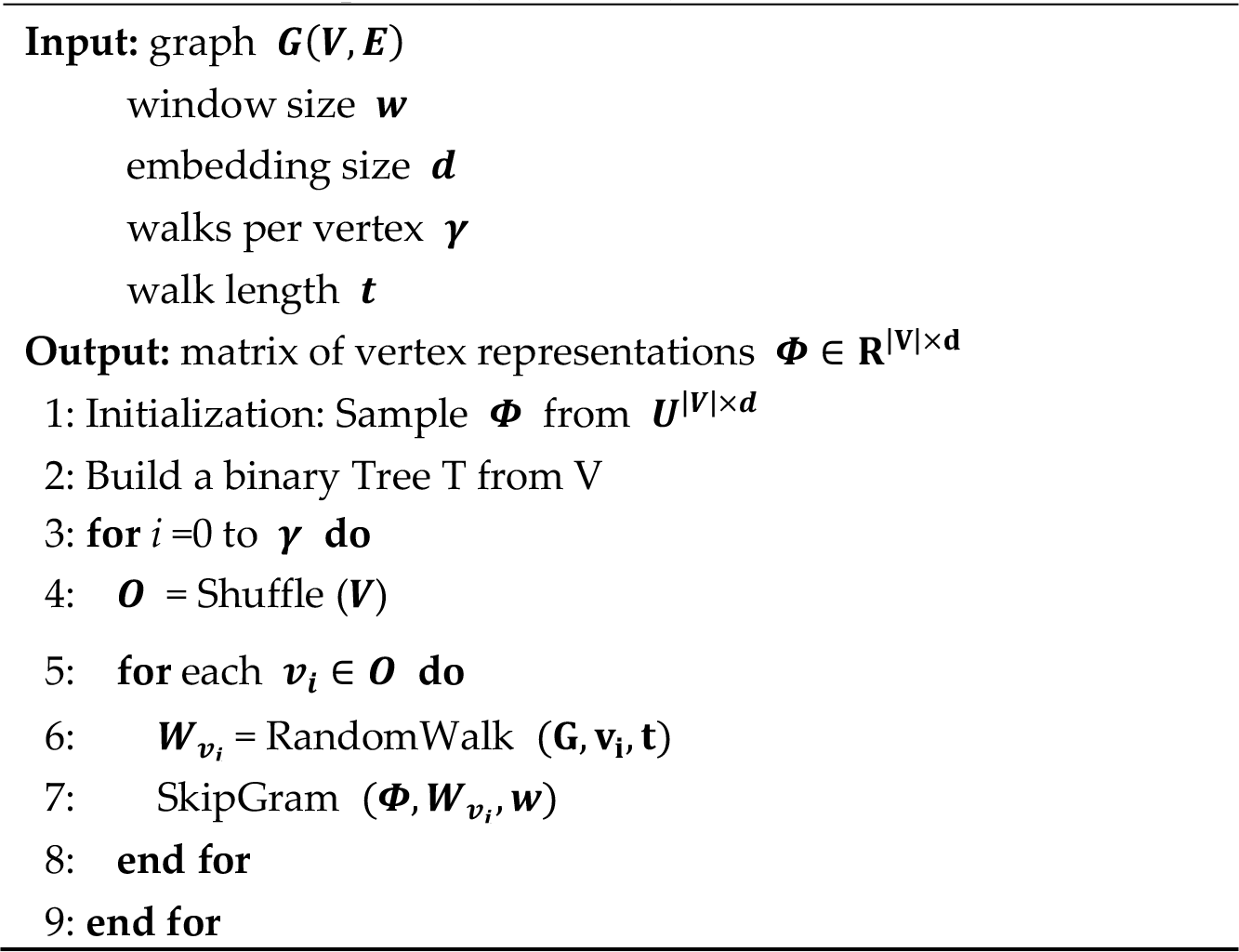

The SkipGram algorithm is as follows:

#### Algorithm 2

SkipGram (***Φ***,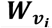,***w***)

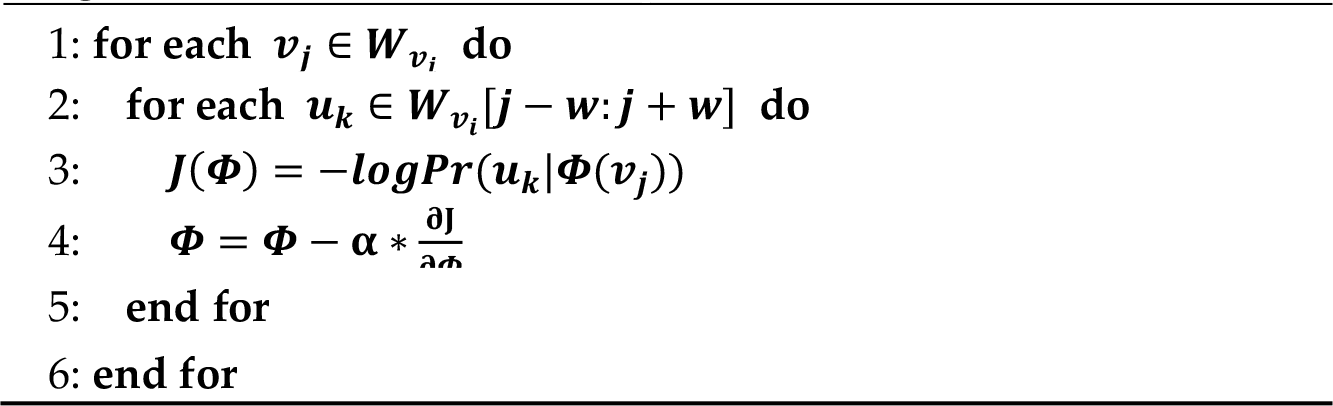

Note whenever the nodes are processed by DeepWalk, the test relationships in the network are stripped to ensure that the label information is not leaked into the test set.

### 2.4 Stack Autoencoder (SAE)

Autoencoder and its variants have been widely used in unsupervised feature learning and classification tasks. Considering that the representation vectors of drug and disease attributes consisting of thousands of dimensions which is not conducive to classifier training. The Stack Autoencoder (SAE) is selected to map vectors of the original space into the new space to reduce noise and make features easy to distinguish. The autoencoder consists of two parts, one is the encoder that maps the original input to the new space, and the other is the decoder that reconstructs the latent representation in the new space back to the original input. For the original input *x*, the output *h*_1_ of the first hidden layer can be calculated by the following formula:

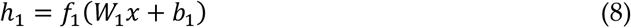

Where *f*_1_ is the activation function, *W*_1_ is the weight matrix between the input layer and the first hidden layer, and *b*_1_ is the threshold of the first hidden layer neurons. Similarly, the output of each layer of the stack autoencoder can be calculated. The mean squared error between the output *y* and the original input *x* is:

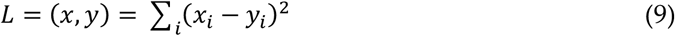

Then the back-propagation algorithm is used to minimize the loss function to get the final model. We completed this task by using the Keras lab in this experiment. The dimension of the hidden layer representation is 64. ‘MSE’ is selected as the loss function and the optimizer is ‘Adam’. The epochs and batch sizes are set to 10 and 128, respectively.

### 2.5 Random Forest Classifier

Random Forest is a classifier that contains multiple decision trees whose output is determined by the mode of the output of each decision tree. It can process high-dimensional features efficiently even in large data volumes. In addition, its high adaptability makes it possible to accept both discrete and continuous data. In this paper, we performed the random forest classifier by a python package called sklearn and all the hyperparameter is set to the default value.

## Results

### 3.1 Relationship prediction based on the whole dataset under 5-fold cross validation

Relationship prediction is a common task in both academia and industry. Here, we will hide a set of edges of the original graph and construct the model based on the incomplete network. Then the hidden edges are utilized for test to assess the proposed method. 5-fold cross validation which is widely used evaluation strategy is applied to carry out this task. In 5-fold cross validation, the whole dataset is divided into 5 mutually exclusive subsets of roughly equal size. Each subset is used as the test set in turn to assess the effect of the classifier, and the remaining 4 subsets are utilized as training set to construct the model. In each fold, area under Receiver Operating Characteristic Curve (ROC) and Precision-Recall Curve (PR) are drawn to visualize the results, respectively. There are total 114,150 experimental valid relationships in the whole network. In each fold cross-validation, 80% edges of the entire network are processed by Biomarker2vec and are treated as the training samples, 20% edges are treated as the test samples.

At the same time, a wide range of evaluation criteria including accuracy (Acc.), sensitivity (Sen.), specificity (Spec.), precision (Prec.) and MCC are adopted to comprehensively and fairly estimate the propose method. The details of results are shown in the following table and figure. Competitive performance under various evaluation criteria demonstrates the keen ability to discover potential associations. The relatively low variance implies the superior robustness and stability of MAN in different situation.

**Table 1.**
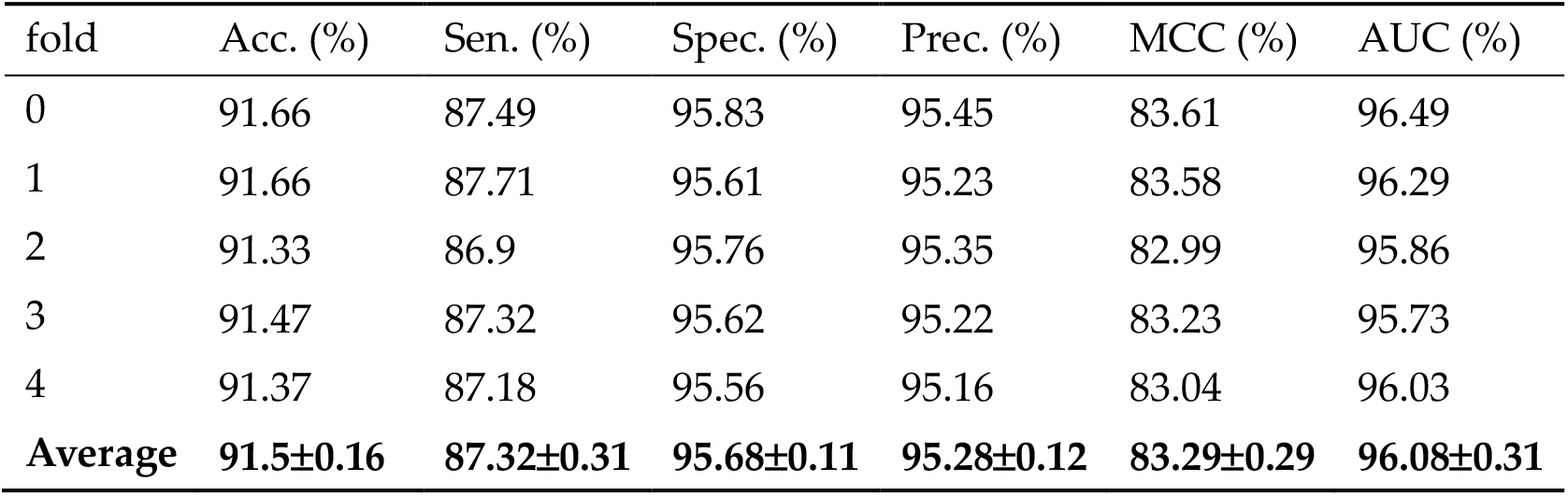
Results of accuracy (Acc.), sensitivity (Sen.), specificity (Spec.), precision (Prec.) and MCC obtained under 5-fold cross validation on the whole network.

**Figure 5.**
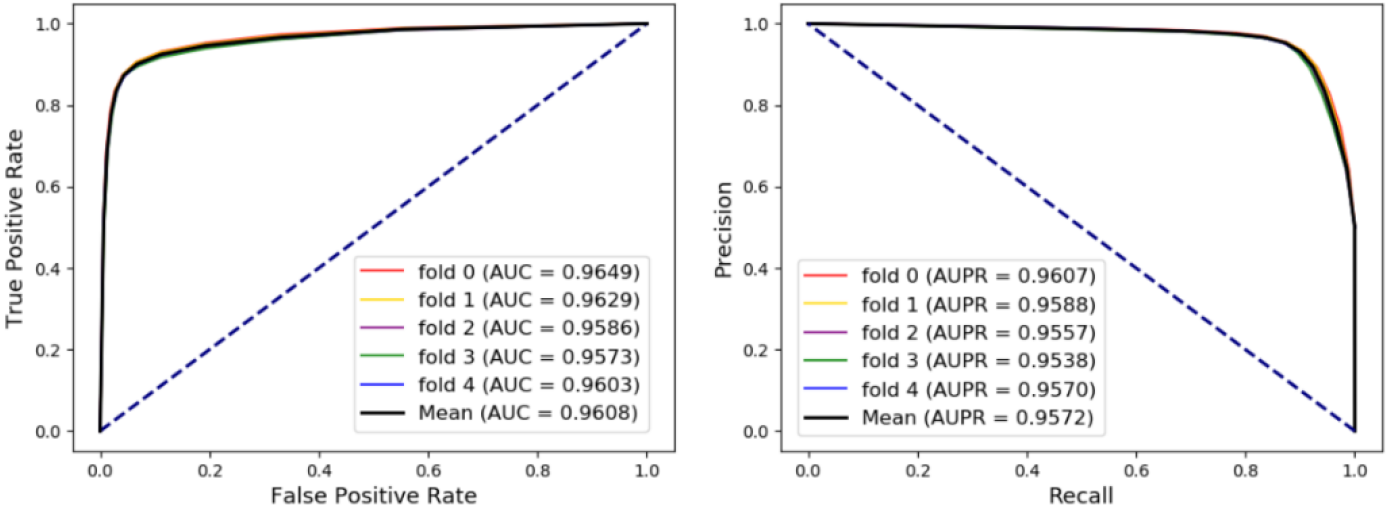
The ROCs, AUCs, PRs and AUPRs obtained under 5-fold cross validation on the whole network

### 3.2 Feature importance comparison

Each node in MAN can be represented as vectors by 2 types of information including node attribute and node behavior. To further evaluate the effectiveness of each kind of feature, we compare the pure attribute-based method, pure behavior-based method and combination of both them based on wide range evaluation metrics, ROC, AUC, PR and AUPR. The results are as in the following table 2 and figure 6.

**Table 2.** Results of accuracy (Acc.), sensitivity (Sen.), specificity (Spec.), precision (Prec.) and MCC obtained by feature importance comparison experiment under 5-fold cross validation on the whole network.

| Feature | Acc. (%) | Sen. (%) | Spec. (%) | Prec. (%) | MCC (%) | AUC (%) |
| --- | --- | --- | --- | --- | --- | --- |
| Attribute | 90.85±0.09 | 89.79±0.19 | 91.9±0.11 | 91.73±0.1 | 81.72±0.17 | 95.91±0.05 |
| Behavior | 88.67±0.15 | 82.15±0.24 | 95.19±0.18 | 94.47±0.19 | 78±0.29 | 93.28±0.13 |
| **Both** | **91.5±0.16** | **87.32±0.31** | **95.68±0.11** | **95.28±0.12** | **83.29±0.29** | **96.08±0.31** |

**Figure 6.**
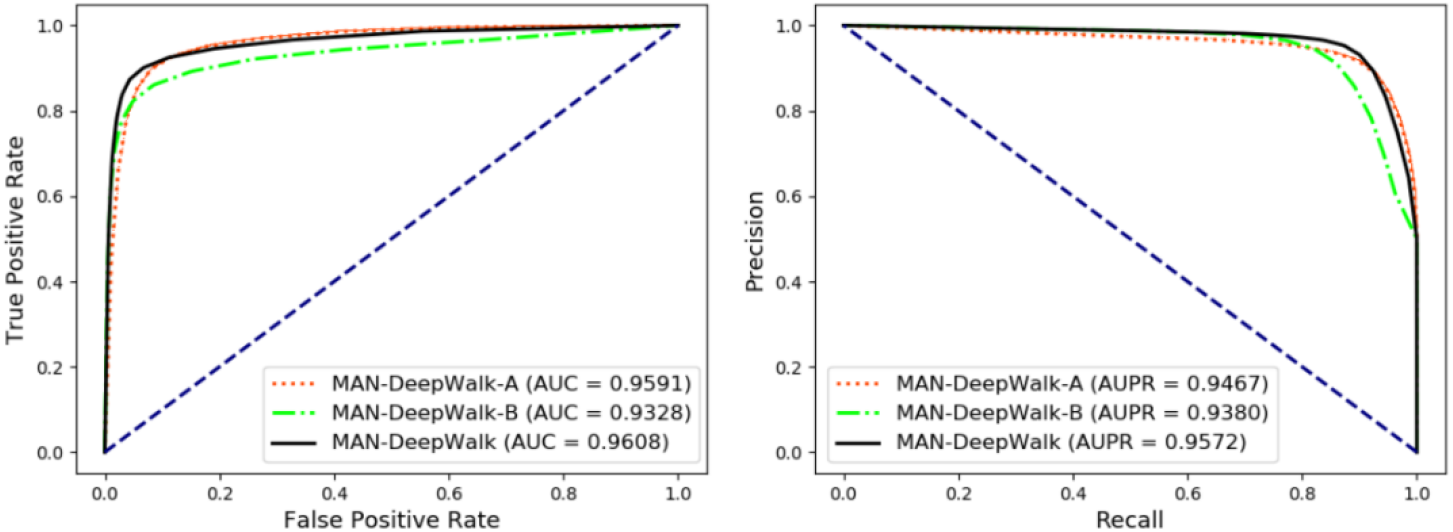
The ROCs, AUCs, PRs, and AUPRs of the proposed method under 5-fold cross validation on the whole dataset.

Based on a single type of feature, the model can achieve considerable prediction performance under 5-fold cross validation, and the more distinguished vectors constructed by combining the above two kinds of information is easier to construct the classifier and achieve more competitive performance.

In view of the “new sample” problem in practical biological experiments, we do not guarantee that the degree of each node is greater than 0. When only the sequences of the biological entities are known and their associations with other biomolecules are undiscovered, this strategy of constructing the vector by combining the node attribute and the node behavior can also predict potential relationships based on new sample and greatly improve the expansion of the model.

### 3.3 Comparison based on varying proportions of training sets

In global relationship prediction, data integrity is considered sensitive and critical. To explore the impact of different ratios of missing data on the results, we separately learn the representation vectors of each node based on varying proportions of edges in the whole graph.

Specifically, 20%, 40%, 60%, and 80% of the edges in the whole network are processed respectively to convert the nodes into vectors by their behavior feature. Meanwhile, the corresponding edges mentioned above are used as the training set to construct the model. The test set is the remaining edges that is 80%, 60%, 40%, and 20% of the whole edges in the graph, respectively. Each node is represented as a vector by only its behavior feature.

Even in the extreme case, *i.e.* 20% of the entire network is used for feature construction and model training, and the remaining 80% edges are used for testing and evaluation. The proposed model still achieved AUC of 0.8710 and AUPR of 0.8747 which implied that the proposed method with outstanding data mining ability will greatly improve the efficiency of existing biological experiments. The details results can be seen in the following table and figure.

**Table 3.**
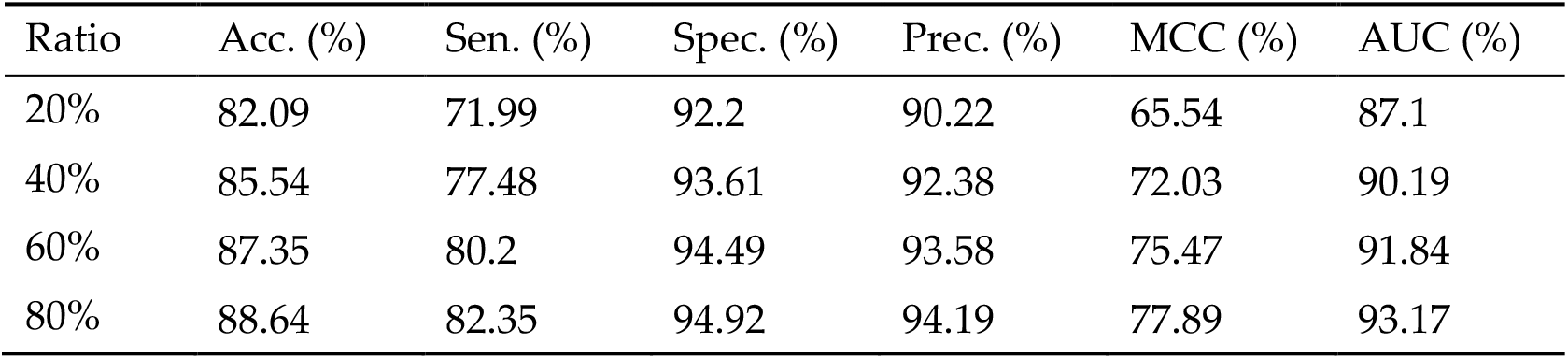
Results of accuracy (Acc.), sensitivity (Sen.), specificity (Spec.), precision (Prec.) and MCC obtained trained and tested by different proportions of edges in the entire network.

| Ratio | Acc. (%) | Sen. (%) | Spec. (%) | Prec. (%) | MCC (%) | AUC (%) |
| --- | --- | --- | --- | --- | --- | --- |
| 20% | 82.09 | 71.99 | 92.2 | 90.22 | 65.54 | 87.1 |
| 40% | 85.54 | 77.48 | 93.61 | 92.38 | 72.03 | 90.19 |
| 60% | 87.35 | 80.2 | 94.49 | 93.58 | 75.47 | 91.84 |
| 80% | 88.64 | 82.35 | 94.92 | 94.19 | 77.89 | 93.17 |

**Figure 7.**
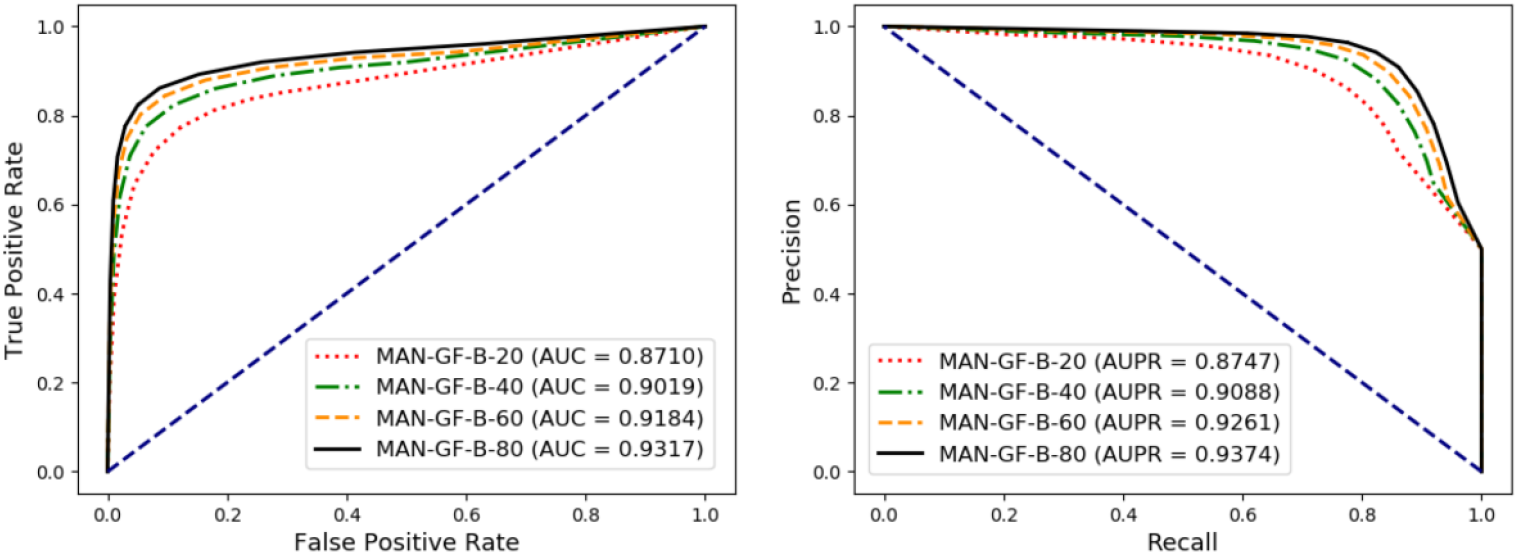
The ROCs, AUCs, PRs, and AUPRs of the proposed method trained and tested by different proportions of edges in the whole network.

### 3.4 Additional experiment based on drug-disease association prediction

Apart from relationship prediction tasks based on the whole network, we take specific object as research subject and further implement additional experiments on drug-disease association prediction to compare the proposed global model with traditional local method. There are 17,414 experimental verified drug-disease associations that have been collected from DrugBank. 5-fold cross validation was performed, and the ROCs and AUCs are shown in the following figure 8.

**Figure 8.**
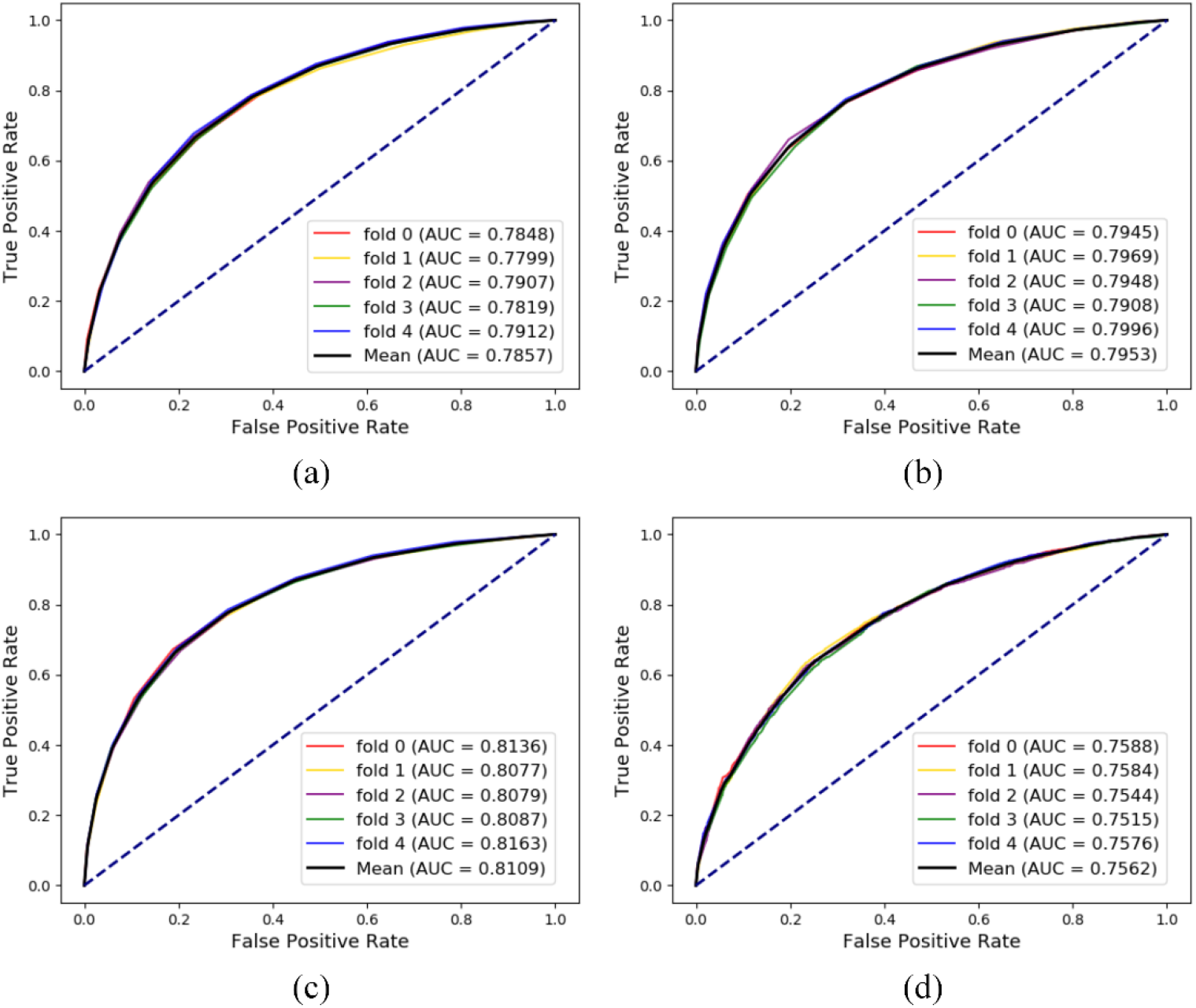
The ROCs, AUCs, PRs, and AUPRs of 4 additional experiments under 5-fold cross validation on dug-disease association dataset.

For figure 8 (a), it can be treated as the baseline that each node is represented as a 64-dimension vector by only its pure attributes *i.e.* Morgan fingerprints or disease semantics.

For figure 8 (b), the node behaviors are represented based on only drug-disease associations. It can be regarded as a traditional idea inspired by the “guilt-by-association” that each node is abstracted into a 128-dimension vector by combining attributes and local behaviors. Compared to figure 8 (a), a slightly elevated AUC confirms the results of the chapter on feature importance comparison and shows that the method of measuring the local function of biomolecules improves the prediction performance to some extent.

For figure 8 (c), it can be considered as a kind of global embedding method that proposed in this paper. In each cross validation, 80% drug-disease pairs along with other all 17 kinds of associations are dealt with Biomarker2vec. Taking the 128-dimension vectors that integrate attribute and behavior as input, Random Forest classifier is chosen for training and testing. The remarkable results compared with traditional local method indicate that the extra edges serve as an intermediary to facilitate the prediction of associations when faced with specific problems.

For figure 8 (d), we carry out a special embedding strategy inspired by Chen *et al.* [53]. The remaining 17 kinds of relationships without drug-disease association pairs are learned by DeepWalk to obtain the behavior representation vectors. Therefore, this process does not depend on any direct drug-disease associations. In order to eliminate the influence of the attribute feature on the prediction performance, each node representation vector was constructed by only behavior feature under the special strategy. Nevertheless, the model still achieved an average AUC of 0.7562 under 5-fold cross validation which implies that MAN does contain a wealth of biological information.

Note that in order to ensure the fairness of the experiment, negative samples of 4 experiments and each subset under 5-fold cross validation are all consistent.

### 3.5 A case study based on drug-disease association

A case study of Ataxia was implemented to assess the performance of the proposed method in a real-world environment. As mentioned above, we have collected 17414 drug-disease associations from CTD database [45] and processed them as described in the article of Zhang *et al.* [54]. In order to verify the predicted effect of the proposed model on new disease, we removed 61 association pairs related to Ataxia, the remaining 17353 drug-disease associations were utilized as the training set to generate feature and construct the model, and each drug is connected to Ataxia in turn to form the test set. The results of top-10 are as follows:

**Table 4.**
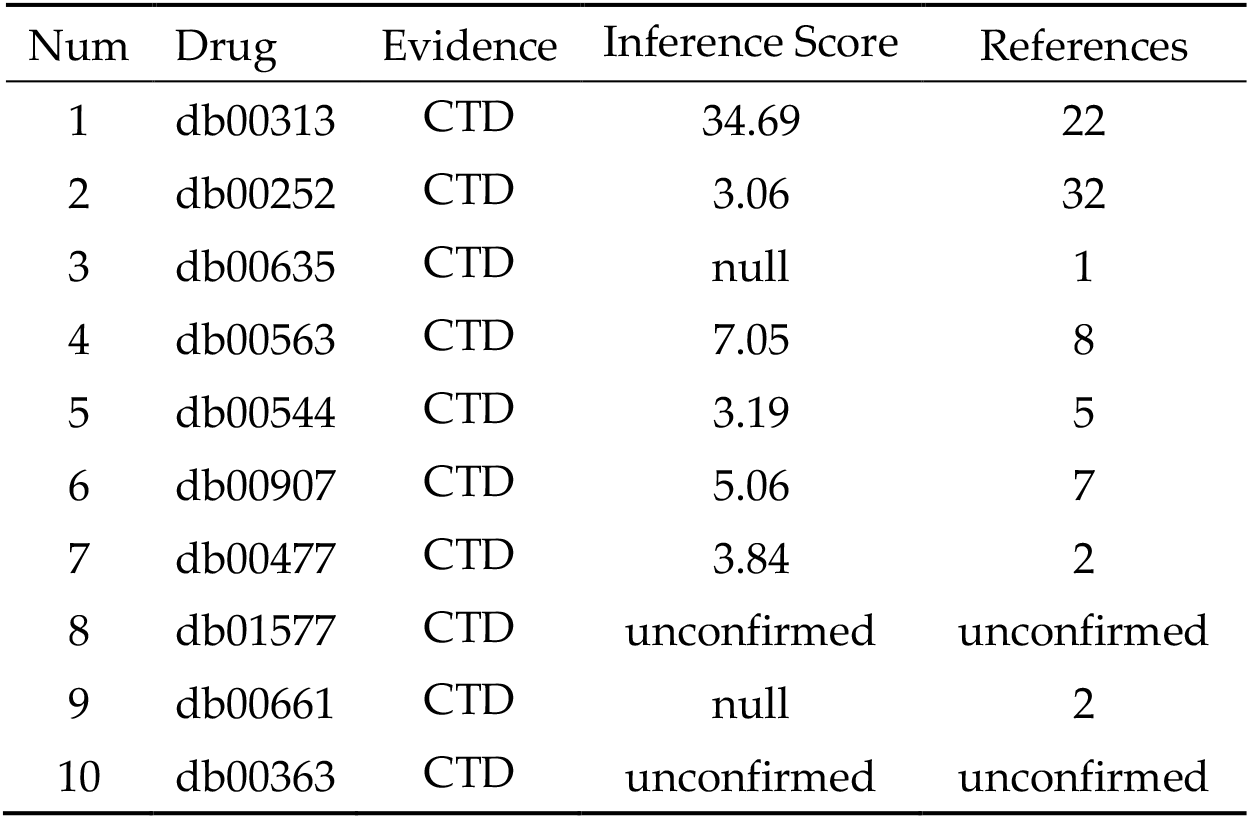
The proposed method was applied to Ataxia to predict the potential disease-related drugs, and 8 of top-10 predicted drugs have been confirmed according to CTD database.

## Conclusion

Current biomarker relationship computational methods can efficiently predict a single logic, but cannot simultaneously detect complex multi-type relationships among various biomarkers. In this article, we abandoned the isolated idea of modeling on only a single range of transcripts or translations, but collected as many kinds of biomolecules as possible to organize a comprehensive Molecular Association Network (MAN). We also proposed a representation method called Biomarker2vec for generating feature vectors of different types of molecules. After above operations, each node (biomarker) in the network (cell) can be characterized as a 128-dimension vector and the random forest classifier is trained to perform relationship prediction task. All results showed that the proposed method achieved remarkable performance in both single-type and multi-type relationship prediction. In general, our studies represent a preliminary exploration from single to complex molecular association network. We believe that this work can bring a technological inspiration and be the basis for further exquisite discovery in alarge practical need. It can be expected there will appear more variants in both proteomics and genomics in the future, from developing new theoretical methods to broader research objects.

## Author Contributions

All authors considered the algorithm, arranged the datasets, and performed the analyses. Z-H.G., Z-H. Y. and Y-B. W. wrote the manuscript. All authors read and approved the final manuscript.

## Funding

This research was funded by the National Natural Science Foundation of China, grant number 61722212, 61902342.

## Conflicts of Interest

The authors declare no conflict of interest.

## References

1. Barabasi A-L, Oltvai ZN: Network biology: understanding the cell’s functional organization. Nature reviews genetics 2004, 5(2):101.

2. Hertzberg RP, Pope AJ: High-throughput screening: new technology for the 21st century. Current opinion in chemical biology 2000, 4(4):445–451.

3. Moore PB: The three-dimensional structure of the ribosome and its components. Annual review of biophysics and biomolecular structure 1998, 27(1):35–58.

4. Mata J, Marguerat S, Bähler J: Post-transcriptional control of gene expression: a genome-wide perspective. Trends in biochemical sciences 2005, 30(9):506–514.

5. Singh R: RNA–protein interactions that regulate pre-mRNA splicing. Gene Expression, The Journal of Liver Research 2002, 10(1-2):79–92.

6. Tian B, Bevilacqua PC, Diegelman-Parente A, Mathews MB: The double-stranded-RNA-binding motif: interference and much more. Nature reviews Molecular cell biology 2004, 5(12):1013.

7. You Z-H, Huang Z-A, Zhu Z, Yan G-Y, Li Z-W, Wen Z, Chen X: PBMDA: A novel and effective path-based computational model for miRNA-disease association prediction. PLoS computational biology 2017, 13(3):e1005455.

8. Guo Z-H, You Z-H, Wang Y-B, Yi H-C, Chen Z-H: A Learning-Based Method for LncRNA-Disease Association Identification Combing Similarity Information and Rotation Forest. iScience 2019, 19:786–795.

9. Wang L, You Z-H, Chen X, Li Y-M, Dong Y-N, Li L-P, Zheng K: LMTRDA: Using logistic model tree to predict MiRNA-disease associations by fusing multi-source information of sequences and similarities. PLoS computational biology 2019, 15(3):e1006865.

10. Li J-Q, You Z-H, Li X, Ming Z, Chen X: PSPEL: in silico prediction of self-interacting proteins from amino acids sequences using ensemble learning. IEEE/ACM Transactions on Computational Biology and Bioinformatics (TCBB) 2017, 14(5):1165–1172.

11. Wang L, You Z-H, Chen X, Yan X, Liu G, Zhang W: Rfdt: A rotation forest-based predictor for predicting drug-target interactions using drug structure and protein sequence information. Current Protein and Peptide Science 2018, 19(5):445–454.

12. Li J-Q, Rong Z-H, Chen X, Yan G-Y, You Z-H: MCMDA: Matrix completion for MiRNA-disease association prediction. Oncotarget 2017, 8(13):21187.

13. Wang Y-B, You Z-H, Li X, Jiang T-H, Chen X, Zhou X, Wang L: Predicting protein–protein interactions from protein sequences by a stacked sparse autoencoder deep neural network. Molecular BioSystems 2017, 13(7):1336–1344.

14. Huang Z-A, Huang Y, You Z-H, Zhu Z, Sun Y: Novel link prediction for large-scale miRNA-lncRNA interaction network in a bipartite graph. BMC medical genomics 2018, 11(6):113.

15. Ashburn TT, Thor KB: Drug repositioning: identifying and developing new uses for existing drugs. Nature reviews Drug discovery 2004, 3(8):673.

16. Chen X, Liu M-X, Cui Q-H, Yan G-Y: Prediction of disease-related interactions between microRNAs and environmental factors based on a semi-supervised classifier. PloS one 2012, 7(8):e43425.

17. Cui H, Zhang M, Yang Q, Li X, Liebman M, Yu Y, Xie L: The Prediction of Drug-Disease Correlation Based on Gene Expression Data. BioMed research international 2018, 2018.

18. Fang S, Zhang L, Guo J, Niu Y, Wu Y, Li H, Zhao L, Li X, Teng X, Sun X: NONCODEV5: a comprehensive annotation database for long non-coding RNAs. Nucleic acids research 2017, 46(D1):D308–D314.

19. Kozomara A, Birgaoanu M, Griffiths-Jones S: miRBase: from microRNA sequences to function. Nucleic acids research 2018, 47(D1):D155–D162.

20. Goyal P, Ferrara E: Graph embedding techniques, applications, and performance: A survey. Knowledge-Based Systems 2018, 151:78–94.

21. Hamilton WL, Ying R, Leskovec J: Representation learning on graphs: Methods and applications. arXiv preprint arXiv:170905584 2017.

22. Wold S, Esbensen K, Geladi P: Principal component analysis. Chemometrics and intelligent laboratory systems 1987, 2(1-3):37–52.

23. Borg I, Groenen P: Modern multidimensional scaling: Theory and applications. Journal of Educational Measurement 2003, 40(3):277–280.

24. Tenenbaum JB, DeSilva V, Langford JC: A global geometric framework for nonlinear dimensionality reduction. science 2000, 290(5500):2319–2323.

25. Roweis ST, Saul LK: Nonlinear dimensionality reduction by locally linear embedding. science 2000, 290(5500):2323–2326.

26. Yao D, Zhang L, Zheng M, Sun X, Lu Y, Liu P: Circ2Disease: a manually curated database of experimentally validated circRNAs in human disease. Scientific reports 2018, 8(1):11018.

27. Zhao Z, Wang K, Wu F, Wang W, Zhang K, Hu H, Liu Y, Jiang T: circRNA disease: a manually curated database of experimentally supported circRNA-disease associations. Cell death & disease 2018, 9(5):475–475.

28. Bao Z, Yang Z, Huang Z, Zhou Y, Cui Q, Dong D: LncRNADisease 2.0: an updated database of long non-coding RNA-associated diseases. Nucleic acids research 2018, 47(D1):D1034–D1037.

29. Fan C, Lei X, Fang Z, Jiang Q, Wu F-X: CircR2Disease: a manually curated database for experimentally supported circular RNAs associated with various diseases. Database 2018, 2018.

30. Bhattacharya A, Cui Y: SomamiR 2.0: a database of cancer somatic mutations altering microRNA–ceRNA interactions. Nucleic acids research 2015, 44(D1):D1005–D1010.

31. Piñero J, Bravo À, Queralt-Rosinach N, Gutiérrez-Sacristán A, Deu-Pons J, Centeno E, García-García J, Sanz F, Furlong LI: DisGeNET: a comprehensive platform integrating information on human disease-associated genes and variants. Nucleic acids research 2016:gkw943.

32. Ma W, Zhang L, Zeng P, Huang C, Li J, Geng B, Yang J, Kong W, Zhou X, Cui Q: An analysis of human microbe–disease associations. Briefings in bioinformatics 2016, 18(1):85–97.

33. Hewett M, Oliver DE, Rubin DL, Easton KL, Stuart JM, Altman RB, Klein TE: PharmGKB: the pharmacogenetics knowledge base. Nucleic acids research 2002, 30(1):163–165.

34. R Rizkallah, Gamal-Eldin S, Saad R, K Aziz R: The pharmacomicrobiomics portal: a database for drug-microbiome interactions. Current Pharmacogenomics and Personalized Medicine (Formerly Current Pharmacogenomics) 2012, 10(3):195–203.

35. Wishart DS, Feunang YD, Guo AC, Lo EJ, Marcu A, Grant JR, Sajed T, Johnson D, Li C, Sayeeda Z: DrugBank 5.0: a major update to the DrugBank database for 2018. Nucleic acids research 2017, 46(D1):D1074–D1082.

36. Chen G, Wang Z, Wang D, Qiu C, Liu M, Chen X, Zhang Q, Yan G, Cui Q: LncRNADisease: a database for long-non-coding RNA-associated diseases. Nucleic acids research 2012, 41(D1):D983–D986.

37. Miao Y-R, Liu W, Zhang Q, Guo A-Y: lncRNASNP2: an updated database of functional SNPs and mutations in human and mouse lncRNAs. Nucleic acids research 2017, 46(D1):D276–D280.

38. Cheng L, Wang P, Tian R, Wang S, Guo Q, Luo M, Zhou W, Liu G, Jiang H, Jiang Q: LncRNA2Target v2. 0: a comprehensive database for target genes of lncRNAs in human and mouse. Nucleic acids research 2018, 47(D1):D140–D144.

39. Yuan J, Wu W, Xie C, Zhao G, Zhao Y, Chen R: NPInter v2. 0: an updated database of ncRNA interactions. Nucleic acids research 2013, 42(D1):D104–D108.

40. Huang Z, Shi J, Gao Y, Cui C, Zhang S, Li J, Zhou Y, Cui Q: HMDD v3. 0: a database for experimentally supported human microRNA–disease associations. Nucleic acids research 2018, 47(D1):D1013–D1017.

41. Liu X, Wang S, Meng F, Wang J, Zhang Y, Dai E, Yu X, Li X, Jiang W: SM2miR: a database of the experimentally validated small molecules’ effects on microRNA expression. Bioinformatics 2012, 29(3):409–411.

42. Chou C-H, Shrestha S, Yang C-D, Chang N-W, Lin Y-L, Liao K-W, Huang W-C, Sun T-H, Tu S-J, Lee W-H: miRTarBase update 2018: a resource for experimentally validated microRNA-target interactions. Nucleic acids research 2017, 46(D1):D296–D302.

43. Tong Z, Cui Q, Wang J, Zhou Y: TransmiR v2. 0: an updated transcription factor-microRNA regulation database. Nucleic acids research 2018, 47(D1):D253–D258.

44. Szklarczyk D, Gable AL, Lyon D, Junge A, Wyder S, Huerta-Cepas J, Simonovic M, Doncheva NT, Morris JH, Bork P: STRING v11: protein–protein association networks with increased coverage, supporting functional discovery in genome-wide experimental datasets. Nucleic acids research 2018, 47(D1):D607–D613.

45. Davis AP, Grondin CJ, Johnson RJ, Sciaky D, McMorran R, Wiegers J, Wiegers TC, Mattingly CJ: The comparative toxicogenomics database: Update 2019. Nucleic acids research 2018, 47(D1):D948–D954.

46. Glažar P, Papavasileiou P, Rajewsky N: circBase: a database for circular RNAs. Rna 2014, 20(11):1666–1670.

47. Shen J, Zhang J, Luo X, Zhu W, Yu K, Chen K, Li Y, Jiang H: Predicting protein–protein interactions based only on sequences information. Proceedings of the National Academy of Sciences 2007, 104(11):4337–4341.

48. Wang D, Wang J, Lu M, Song F, Cui Q: Inferring the human microRNA functional similarity and functional network based on microRNA-associated diseases. Bioinformatics 2010, 26(13):1644–1650.

49. Weininger D: SMILES, a chemical language and information system. 1. Introduction to methodology and encoding rules. Journal of chemical information and computer sciences 1988, 28(1):31–36.

50. Rogers D, Hahn M: Extended-connectivity fingerprints. Journal of chemical information and modeling 2010, 50(5):742–754.

51. Landrum G: RDKit: open-source cheminformatics software. In.; 2016.

52. Perozzi B, Al-Rfou R, Skiena S: Deepwalk: Online learning of social representations. In: Proceedings of the 20th ACM SIGKDD international conference on Knowledge discovery and data mining: 2014: ACM; 2014: 701–710.

53. Chen X: Predicting lncRNA-disease associations and constructing lncRNA functional similarity network based on the information of miRNA. Scientific reports 2015, 5:13186.

54. Zhang W, Yue X, Lin W, Wu W, Liu R, Huang F, Liu F: Predicting drug-disease associations by using similarity constrained matrix factorization. BMC bioinformatics 2018, 19(1):233.

